# Cross-species metabolomic analysis of DDT and Alzheimer’s disease-associated tau toxicity

**DOI:** 10.1101/2021.06.14.448355

**Authors:** Vrinda Kalia, Megan M. Niedzwiecki, Joshua M. Bradner, Fion K. Lau, Meghan L. Bucher, Katherine E. Manz, Zoe Coates Fuentes, Kurt D. Pennell, Martin Picard, Douglas I. Walker, William T. Hu, Dean P. Jones, Gary W. Miller

## Abstract

**Background:** The formation of hyperphosphorylated tau (p-tau) protein tangles in neurons is a pathological marker of Alzheimer’s disease (AD). Exposure to the pesticide dichlorodiphenyltrichloroethane (DDT) has been associated with increased risk of AD.

**Objectives:** To determine if there was a connection between DDT exposure and tau toxicity we investigated whether exposure to DDT can exacerbate tau protein toxicity in *C. elegans.* In addition, we examined the association between p-tau protein and metabolism in a human population study and in a transgenic *C. elegans* strain neuronally expressing a mutant tau protein fragment that is prone to aggregation.

**Methods:** In the human population study, we used a metabolome-wide association framework to determine the association between p-tau measured in the cerebrospinal fluid (CSF) and metabolomic features measured in both plasma (n = 142) and CSF (n = 78) using high-resolution metabolomics (HRM). Using the same HRM method, we determined changes in metabolomic features in the transgenic *C. elegans* strain compared to its control strain. Metabolites associated with p-tau in both species were analyzed for overlap. We also examined the effect of DDT and aggregating tau protein on growth, swim behavior, mitochondrial function, metabolism, learning, and lifespan in *C. elegans*.

**Results:** Plasma and CSF-derived features associated with p-tau level were related to drug, amino acid, fatty acid and mitochondrial metabolism pathways. Five metabolites overlapped between plasma and *C. elegans*, and 4 between CSF and *C. elegans*. DDT exacerbated the inhibitory effect of aggregating tau protein on growth and basal respiration. In the presence of aggregating tau protein, DDT induced more curling and was associated with reduced levels of amino acids but increased levels of uric acid and adenosylselenohomocysteine. Developmental exposure to DDT blunted the lifespan reduction caused by aggregating tau protein.

**Conclusion:** The model organism *C. elegans* can complement human studies by providing a means to study mechanisms of environmental toxicants. Specifically, our *C. elegans* data show that DDT exposure and tau protein aggregation both inhibit mitochondrial function and DDT exposure can exacerbate the mitochondrial inhibitory effects of tau protein aggregation providing a plausible explanation for the observed human associations.

## Introduction

In 2014, 5 million people in the US were living with Alzheimer’s disease (AD). By 2060, this number is projected to grow to 13.9 million (Matthews et al. 2019). Clinically, AD manifests as dementia, a progressive deterioration of memory and cognitive function (Van Cauwenberghe et al. 2016). Pathologically, AD is characterized by severe neuronal loss, aggregation of amyloid-β (Aβ) in extracellular senile plaques, and formation of intraneuronal neurofibrillary tangles consisting of hyperphosphorylated tau (p-tau) protein. There is evidence that AD may be a metabolic neurodegenerative disease (de la Monte and Wands 2008) as it has been associated with altered local and peripheral metabolism in several studies. Through *in vitro*, animal model, and epidemiological studies, investigators have found associations between tau neurofibrillary tangles and impaired glucose metabolism (Bischof and Park 2015; Ossenkoppele et al. 2015, 2016), altered mitochondrial trafficking, morphology and bioenergetics, and reduced ATP production (Pérez et al. 2018). While aging is the strongest risk factor of AD, evidence of risk factors for dementias show that lifestyle choices and the environment may modify disease onset and alter the projected prevalence (Nichols et al. 2019). Indeed, using untargeted high-resolution metabolomics (HRM), our group has uncovered plasma derived metabolites from endogenous and exogenous sources associated with the disease (Niedzwiecki et al. 2020; Vardarajan et al. 2020). If causally associated with the disease, these metabolites may be modified or targeted to alter disease prevalence or progression.

The role of the environment in AD pathogenesis has a controversial history (DeKosky and Gandy 2014) but recent studies provide evidence of environmental chemical exposures influencing disease risk. In a cross-sectional, case-control study, Richardson and colleagues found that cases of AD had higher levels of a metabolite of the pesticide DDT (1,1,1-tricholoro-2,2-bis(p-chloro-phenyl) ethane) in their serum (Richardson et al. 2014). DDT is a highly persistent, synthetic organochlorine pesticide used for pest control in agricultural settings and to control vectors that can cause diseases like malaria and typhus. It was widely used in the USA from 1939 to 1972, until its use was banned by the US Environmental Protection Agency (EPA) (Turusov et al. 2002). Despite its regulation, DDT and its metabolites remain persistent and can be detected in the blood of most of the US population (National Report on Human Exposure to Environmental Chemicals | CDC 2020). Additionally, DDT can be passed through breastmilk to infants, exposing generations that have been born after its ban (Needham et al. 2011). DDT is still used for vector control in some African and south Asian countries (van den Berg et al. 2017) and can travel long distances through evaporation, distillation, and transport via winds and ocean currents (Wania and Mackay 1996). Therefore, DDT poses a threat to the health of populations living in countries where it is still produced and in countries that are further away.

*Caenorhabtidis elegans* (*C. elegans* or worms) is a non-parasitic nematode that has long been used in neuroscience and developmental research; more recently it has been gaining popularity as an *in vitro* model in toxicity testing. Studies in *C. elegans* show that the toxicity ranking of several toxicants, including, but not limited to, metals, organophosphate pesticides, ∼ 60% of chemicals in the EPA’s ToxCast™ Phase I and Phase II libraries, known or suspected developmental toxicants, and metabolic toxicants, is predictive of rat LD_50_ values (Boyd et al. 2016; Cole et al. 2004; Harlow et al. 2016; Hunt 2017; Hunt et al. 2012; Middendorf and Dusenbery 1993; Williams and Dusenbery 1988). The model is inexpensive and requires minimal laboratory expertise to maintain. Several fundamental aspects of biology were discovered in *C. elegans* including apoptosis, RNAi, and miRNA. Furthermore, *C. elegans* are the first complex organism to have their genome sequenced (*C. elegans* Sequencing Consortium 1998), allowing access to a large library of genetic mutant strains. The long history of its use in biology and the conservation of several genes and pathways between worms and humans (Kaletta and Hengartner 2006) makes the nematode model valuable for biological insight (Brenner 1974; Corsi et al. 2015), particularly to study gene-environment interactions. In this study, we use untargeted liquid chromatography (LC) with high-resolution mass spectrometry to identify plasma and cerebrospinal fluid (CSF) derived metabolites associated with p-tau levels measured in the CSF of individuals from a clinical study of AD. We then compared metabolites associated with p-tau to the metabolic profile of a transgenic strain of *C. elegans* that is a model of AD-related pathology which expresses a mutant fragment of tau protein in all neurons (Fatouros et al. 2012). Using the same mutant tau transgenic strain of *C. elegans* (Fatouros et al. 2012), we then tested the effect of exposure to DDT on growth, behavior, metabolism, learning, and survival.

## Methods

### 1. Chemicals

BD Bacto dehydrated agar, salts to make M9 buffer (monobasic potassium phosphate, dibasic sodium phosphate, sodium chloride, magnesium sulfate), 1N sodium hydroxide, p,p’-DDT (>95%), dimethyl sulfoxide (DMSO, 99.9%), acetone (99.8%, HPLC grade), n-hexane (>=99%), dichloromethane (99.8%, HPLC grade), sodium azide (99%), and 2-butanone (≥ 99.0%) were purchased from Fisher Scientific (Waltham, MA). Carbonyl cyanide 4-(trifluoromethoxy) phenylhydrazone (FCCP, > 98%) was purchased from Sigma-Aldrich (St. Louis, MO). Certified reference standards for GC-HRMS quantification were purchased from Accustandard (New Haven, CT), including o,p’-DDT, p,p’-DDT, o,p’-DDE, p,p’-DDE, ^13^C_12_ labeled p,p’-DDE, D_8_ labeled p,p’-DDT, phenanthrene D-10, and chrysene D-12. The hypochlorite solution used for synchronization was prepared using household bleach (Clorox, 8% sodium hypochlorite), water, and 1N sodium hydroxide.

### 2. Human: participants and sample collection

This study was approved by the Emory University Institutional Review Board and the methods have been previously described (Niedzwiecki et al. 2020). Briefly, subjects were recruited from the Emory Cognitive Neurology Clinic and the Emory Alzheimer’s Disease Research Center. Each subject underwent a detailed neurological and neuropsychological evaluation. Subjects were classified as having normal cognition (NC) if there was no subjective cognitive complaint and neuropsychological analysis showed normal cognitive functioning for their age, gender, education, and race; mild cognitive impairment (MCI) (Albert et al. 2011), or AD dementia (McKhann et al. 2011) according to NIA-AA criteria (Niedzwiecki et al. 2020). Plasma and CSF samples were collected and processed as described previously (Hu et al. 2015; Niedzwiecki et al. 2020). CSF AD biomarker analysis was performed as previously described using a Luminex 200 platform to determine levels of total tau (t-Tau), and tau phosphorylated at threonine 181 (p-Tau_181_) (Howell et al. 2017).

### 3. C. elegans methods

#### 3.1. *C. elegans*: growth and maintenance

Standard methods of culture, including the use of normal or high growth media (NGM/HGM) plates, culture temperature of 20 °C and the OP50 *E. coli* strain as a food source, were followed as described (Brenner 1974) unless noted otherwise. *C. elegans* strains used included the wild type N2 Bristol strain, BR5271 (*byIs162* [P*_rab-3_*::F3(delta)K280 I277P I380P + P*_myo-2_*::mCherry]; referred to as the “non-aggregating/non-agg” strain), and BR5270 (*byIs161* [P*_rab-3_*::F3(delta)K280 + P*_myo-2_*::mCherry]; referred to as the “aggregating/agg” strain). All strains were provided by the Caenorhabditis Genetics Center, which is funded by NIH Office of Research Infrastructure Programs (P40 OD010440).

#### 3.2. *C. elegans*: exposure to DDT

Worms were exposed to the pesticide p,p’-DDT or the solvent control, DMSO, on NGM plates. DDT exposure plates were created using methods previously described (Hunt et al. 2011). Briefly, a 20 mM stock of DDT, made by dissolving in 100% DMSO, was diluted to 150 μM with sterile water and then applied on the surface of NGM plates spotted with OP50 *E. coli* to obtain the appropriate final concentration of DDT on the plate. The solvent control plates were created following the same dilution but without DDT to achieve a final concentration of 0.015% DMSO. DDT was allowed to diffuse and the plates were allowed to equilibrate for 2 hours before worms were introduced. All worms were exposed to a final concentration of 3 μM DDT unless otherwise stated.

#### 3.3. *C. elegans*: DDT uptake experiments

A synchronized population of wildtype worms, created using hypochlorite treatment, were grown on 10 cm NGM plates with 0.3, 3, or 30 μM DDT, and the DMSO control. The non-aggregating and aggregating worms were similarly synchronized and exposed to 3 μM DDT or DMSO. All strains were collected after 72 hours of exposure, at the young adult stage. They were washed in M9 buffer 4x and sorted into aliquots of 1000-1200 worms using the COPAS FP-250. The volume of M9 buffer in each sample was reduced to 100 μL and each sample was snap frozen in liquid nitrogen. To extract DDT and its metabolites, the worm cuticle was disrupted by bead beating (6.5 m/s for 1 minute) and the samples were analyzed for levels of DDT.

#### 3.4. *C. elegans*: growth determined through size measured on COPAS Biosorter

The COPAS Flow Pilot (FP) 250 is an instrument used for high-throughput manipulation of *C. elegans*. For each worm that passes through the flow cell, the COPAS FP-250 determines its time-of-flight (TOF), which represents the length of the worm passing through the flow cell, and the extinction, which represents the optical density or thickness of the worm passing through the flow cell. After hypochlorite synchronization, eggs from all three strains were allowed to hatch and develop on 10 cm NGM plates with 3 μM DDT or DMSO. Worms were sorted through the COPAS FP 250 to measure TOF and extinction 46-50 hours after synchronization (around the L4 stage, *n* = 1000-4000 per group, object inclusion criteria: Log(TOF) > 6 and Log(extinction) > 5) and at 70-72 hours post synchronization (young adults, *n* = 1000-4000 per group, object inclusion criteria: Log(extinction) between 5.5 and 9). We used inclusion criteria that have been previously estimated for the L4 larval and adult stage on the COPAS FP-250 (Boyd et al. 2016). Measures of TOF and extinction were compared across the strains and treatment groups using a one-way analysis of variance at the two time points.

#### 3.5. *C. elegans*: swim behavior

The celeST software package was used to determine aspects of swim behavior for the different strains exposed to DDT or the solvent control (Restif et al. 2014). Briefly, 3-4 worms at the young adult stage were placed in 60 μL of M9 buffer in a 15 mm ring preprinted on a microscope slide (Fisherbrand microscope slides with two 15 mm diameter circles, catalog #22-339-408). Recordings of swim behavior were made as a series of jpeg images using a chameleon 3 camera (FLIR, Wilsonville, OR) for 30 s at a frame rate of 18 f/s. Data were collected from four-five trials representing different experiments, with a total of 50-100 worms recorded per group.

#### 3.6. *C. elegans*: seahorse XFe96 extracellular flux analysis

The three strains exposed to DDT or solvent control were collected at the young adult stage and washed in M9 buffer 4x for analysis using the Seahorse XFe96 extracellular flux analyzer (Koopman et al. 2016). Briefly, 3-30 worms in M9 buffer were plated into the wells of a Seahorse utility plate and the volume of M9 buffer in each well was made up to 200 μL. M9 buffer without any worms was used as the blank for background correction. Baseline respiration was measured (measurement numbers 1-5), followed by injection of FCCP (10 μM, final concentration) to elicit maximal respiration (measurement number 6-14), followed by sodium azide (40 mM, final concentration) to measure non-mitochondrial respiration (measurement number 15-18). Data were normalized to the number of worms in each well to determine the rate of oxygen consumption (pmol O_2_/min) per worm. Basal respiration was determined as the difference between non-mitochondrial respiration and the average oxygen consumption rate at measurements 2 through 5; maximal respiration was determined as the difference between non-mitochondrial respiration and the average oxygen consumption rate measured after the FCCP injection; and spare respiratory capacity was measured as the difference between basal and maximal respiration.

#### 3.7. *C. elegans*: associative learning assay

The associative learning assay was carried out as previously described (Kauffman et al. 2011) with some modifications. The assay relies on an associative memory paradigm where worms are trained by pairing the presence of food with the odor of 10% butanone. Briefly, worms were hypochlorite synced and allowed to grow on DDT or solvent control plates for about 72 hours, until they reached the young adult stage. Worms were collected off plates and washed 3x in M9 buffer. After the last wash, the naïve attraction toward butanone was assessed. Worms were then starved for an hour, after which the conditioned training was performed to pair the odor with the presence of food. The attraction toward butanone was determined just after conditioning, representing their ability to learn and form an associative memory. To count the number of worms attracted to the butanone spot or the control (95% ethanol) spot, images of the entire assay plates were taken on a Basler GigE camera, and the images were analyzed using a MATLAB algorithm created by the Murphy lab (Kauffman et al. 2011).

#### 3.8. *C. elegans*: survival analysis

Wildtype worms and the transgenic strains were exposed to DDT or DMSO until the young adult stage, around 72 hours. After exposure, worms were collected off plates and washed in M9 buffer 4 times. We created 4 replicates per treatment group per strain with 25-35 worms each. Adult worms were counted and transferred everyday onto new 6 cm NGM plates until they stopped producing progeny (∼ adult day 6). At this point, worms were transferred onto 6 cm NGM plates with nystatin and ampicillin. Worms were then counted every other day and scored as dead if they did not respond to the gentle touch of a platinum wire. A worm was censored from the plate if it was missing, showed internal hatching, or was damaged during transfer. Worms were followed until they were all dead. Data was analyzed using Kaplan Meier survival calculations and the log-rank test using the R package survival.

### 4. High-resolution mass spectrometry methods

#### 4.1. Gas chromatography with high-resolution mass spectrometry (GC-HRMS) for *C.* elegans DDT uptake

Worm tissue concentrations of p,p’-DDT, p,p’-DDE, p,p’-DDD, o,p’-DDT, o,p’-DDE and o,p’-DDD were measured using methods previously described (Elmore et al. 2020). Prior to extraction, each sample was spiked with labeled isotope internal standards to assess analyte recovery. Each sample was extracted using QuEChERS (Quick, Easy, Cheap, Effective, Rugged, Safe). The samples were first vortex mixed in centrifuge tubes with 1 mL 1:1:1 hexane:acetone:dichloromethane and sonicated for 30 minutes. The entire sample and the supernatant were transferred to a centrifuge tube containing 150 g MgSO4 and 50 mg C18 (United Chemical Technologies, Bristol, PA), vortexed for 30 seconds, and centrifuged for 5 minutes at 1,105 x g. This extraction was repeated 2 more times and the final 3 mL extract was evaporated to 150 μL under nitrogen (Organomation 30 position Multivap Nitrogen Evaporator), transferred to a low-volume (300 μL) GC vial, and spiked with phenanthrene-D10 and chrysene-D12 as volumetric internal standards to ensure injection consistency during GC-HRMS analysis. Extracts were analyzed on a GC Q-Exactive Orbitrap MS (Thermo Scientific) equipped with a Thermo Trace 1300 gas chromatograph and TriPlus RSH Autosampler using chromatographic methods described previously (Elmore et al. 2020). The MS was operated in full scan mode, with a scan range of 50 to 750 m/z. Analytes were quantified using the most abundant fragment and identity was confirmed using the ratio of two confirming ions and retention times (Supplemental Table 1).

#### 4.2. Sample preparation for liquid chromatography coupled high resolution mass spectrometry (LC-HRMS)

Human sample preparation: Samples were prepared for HRM using methods detailed elsewhere (Go et al. 2015; Niedzwiecki et al. 2020; Park et al. 2012; Soltow et al. 2013). Briefly, aliquots of plasma or CSF were removed from −80 °C storage and thawed on ice. 65 μL of each biofluid was added to 130 μL of acetonitrile containing a mixture of stable isotopic standards, vortexed, and allowed to equilibrate for 30 min. Proteins were precipitated by centrifuge (16,100 x g at 4 °C for 10 min) and the supernatant was transferred to a low-volume vial for analysis.

*C. elegans* sample preparation: two different experiments were conducted and prepared: 1. To determine the metabolic effects of aggregating tau protein, a synchronized population of worm eggs of the non-aggregating and aggregating strain were allowed to hatch and grow on NGM plates. 2. To determine the effect of DDT on metabolism in all strains, a synchronized population of all strains, wildtype, non-aggregating and aggregating worms, were placed on NGM plates coated with DDT or DMSO. In both experiments, worms were allowed to grow until larval stage 4. For collection, worms were washed 4x in M9 buffer and sorted into four-to-six replicates containing 500 worms using a COPAS FP-250. The final volume was reduced to 100 μL by centrifuge and each sample was snap frozen in liquid nitrogen and stored at −80 °C until needed for processing. Metabolites were extracted using methods described previously (Bradner et al. 2021; Mor et al. 2020). Briefly, two volumes of acetonitrile (200 μL) containing a mixture of internal standards was added to the 100μL worm suspension, and samples were homogenized by bead-beating. A spatula-full of zirconium oxide beads (∼10 beads, 0.5 mm diameter, Yttria stabilized) from Next Advance (Troy, NY) was added to each worm sample, and placed in a bead beater (Next Advance Bullet Blender Storm, Troy, NY) set at 6.5 m/s for 30 seconds. Extracts were then allowed to equilibrate on ice for one minute, and placed in the beater for another 30 seconds at the same speed. After equilibration on ice for 30 minutes, proteins were removed by centrifuge (15,000 x g at 4 °C for 10 min). All sample processing was performed on ice or in a cold room when necessary.

#### 4.3. High-resolution metabolomic analyses

Human sample extracts were analyzed by reverse-phase C18 liquid chromatography (Dionex Ultimate 3000) and Fourier transform mass spectrometry in positive electrospray ionization mode, resolution (FWHM) of 70,000 (Niedzwiecki et al. 2020). Sample extracts from *C. elegans* were analyzed on an LC-HRMS platform in two ways: 1. For determination of the effect of aggregating tau protein on metabolism, sample extracts were analyzed using untargeted LC-HRMS using methods described previously (Liu et al. 2020). Mass spectral data were generated under positive electron spray ionization in full scan mode. 2. Due to changes in LC-HRMS technologies, analysis of DDT exposure studies used slightly different analytical conditions; however, detection of endogenous metabolites across the two platforms is consistent. For determination of the effect of DDT on metabolism in all strains, after processing, the supernatant was diluted 1:1 in HPLC-grade water and analyzed using a HILIC column (positive and negative ESI mode) and C18 column (positive and negative ESI mode). Separation was similar to conditions described above, except an acetonitrile gradient with 10 mM ammonium acetate was used for HILIC, and acetonitrile gradient with 0.5% acetic acid for C_18_. For both methods, 10 μL of the sample extract was injected in triplicate. All mass spectral data were generated on an orbitrap mass spectrometer in full scan mode (1: Thermo Scientific Q-Exactive HF and 2: Thermo Scientific HFX), scanning for mass range 85 to 1250 Da. All raw mass spectral data were extracted using the R packages apLCMS (Yu et al. 2009) and xMSanalyzer (Uppal et al. 2013). Due to the need for multiple batches in the human study, batch correction was performed using ComBat (Leek et al. 2012). No batch effects were observed for *C. elegans* studies and detected intensities were used as is for statistical analyses. Intensities were generalized log transformed prior to analysis.

### 5. High-resolution metabolomic data analyses

#### 5.1. Human: analysis of LC-HRMS data

Association of metabolite peaks with p-tau levels were assessed using linear regression for metabolites detected in >80% of study samples while controlling for sex, age and analysis batch.

Features associated with p-tau levels (*p* < 0.05) were analyzed for metabolic pathway enrichment using mummichog (version 2.0.6) in Python (version 2.7) (Li et al. 2013).

In a sensitivity analysis, features associated with AD dementia vs. NC were examined using the subset of features present in >20% of samples, reflecting a markedly lower threshold for feature filtering compared to our previous analysis of AD (Niedzwiecki et al. 2020). Statistical analyses were conducted as previously described (Niedzwiecki et al. 2020).

#### 5.2. *C. elegans*: analysis of LC-HRMS data and overlap

All feature tables were processed as follows: first, the intensity of a metabolite peak in the samples was compared to its intensity in the medium blank (M9 buffer). If the intensity was ≥ 1.5 times the intensity in the blank in all samples, it was retained for subsequent analysis. Second, if a metabolite peak was missing from fewer than 50% of the samples, it was replaced with half the value of the minimum intensity measured in the samples. Features missing from more than 50% of the samples were removed from downstream analysis. Third, the filtered and imputed feature table was imported into MetaboAnalyst (Pang et al. 2020) and normalized by generalized log transformation. Three different analyses were conducted using the processed *C. elegans* data:

1. Metabolic effects of aggregating tau protein. The filtered feature table was used to determine the metabolites associated with the aggregating worms by comparing aggregating to non-aggregating worms using multiple t-tests. Metabolites with p < 0.05 were analyzed for pathway analysis using mummichog hosted on MetaboAnalyst (Chong et al. 2018) using the *C. elegans* KEGG reference map.
2. Analysis of metabolites common to humans and *C. elegans* that are associated with tau protein. Plasma and CSF derived metabolites that were associated with CSF p-tau (*p* < 0.05) or neuronal expression in *C. elegans* (*p* < 0.05) were compared using KEGG ID annotations from pathway analysis. A metabolite was considered overlapping if it was significantly associated with tau protein in both species, was annotated with a KEGG ID, and the direction of association was concordant between worms and humans.
3. The metabolic effect of DDT in all three strains. The filtered feature table was used to determine: the metabolomic profile of DDT exposure in wildtype worms, the metabolomic profile of DDT exposure in the aggregating worms, and the metabolomic profile associated with the aggregating worms by comparing metabolite peaks from the aggregating and non-aggregating worms exposed to the vehicle control. For all analyses, we used t-tests to compare differences in mean intensities. Significant metabolite peaks were tested for pathway enrichment using mummichog on MetaboAnalyst and the *C. elegans* KEGG reference map.

### 6. Statistical analyses

Tests for significance were determined through one-way ANOVA and post-hoc Tukey’s HSD test, unless stated otherwise. All data were analyzed in R (version 4.0.2) using RStudio (v1.1.456) unless otherwise stated. Code and data associated with worm assays can be found at: https://github.com/vrindakalia/DDT_tau_Celegans

## Results

### CSF and plasma metabolism associated with CSF p-tau levels

There were no significant differences in the distribution of age or gender between the three different diagnoses. Patients with AD and MCI had higher levels of p-tau measured in their CSF compared to controls (Table 1). Following data extraction and filtering, 6,028 metabolite peaks were detected and measured in the CSF of patients and 7,249 m/z features in plasma. After controlling for the age, sex, and batch of analysis, we found 225 metabolites in CSF (Figure 1 A) and 391 in plasma (Figure 1 C) that were associated with CSF levels of p-tau at *p* < 0.05. Pathway analysis of the CSF metabolites found pathways associated with glutamate metabolism, carnitine metabolism, lysine metabolism, saturated fatty acid metabolism, as well as metabolism of several amino acids (Figure 1 B), while pathways in plasma-derived were consistent with drug metabolism, carnitine metabolism, lysine metabolism, and pathways associated with energy production (Figure 1 D).

**Table 1.** Demographic data of patients. Patients with a diagnosis of Alzheimer’s disease (AD) or mild cognitive impairment (MCI) were included in the analysis. Of these patients, 142 plasma and 78 cerebrospinal fluid (CSF) samples were analyzed using the LC-HRMS method. t-Tau: total tau protein; p-Tau181: tau phosphorylated at threonine 181.

|  | Control | AD | MCI |
| --- | --- | --- | --- |
| **PLASMA (N)** | 46 | 51 | 45 |
| % Male | 30 | 35 | 48 |
| Age (y, mean ± SD) | 66.5 ± 8.7 | 65.9 ± 8.9 | 69.4 ± 6.6 |
| CSF t-Tau (pg/ml, mean ± SD) | 44 ± 23 | 120 ± 66 | 76 ± 70 |
| CSF p-Tau181 (pg/ml, mean ± SD) | 32 ± 14 | 74 ± 30 | 51 ± 25 |
| **CSF (N)** | 25 | 26 | 27 |
| % Male | 28 | 38 | 55 |
| Age (y, mean ± SD) | 66.2 ± 8.2 | 64.8 ± 8.2 | 70.2 ± 6.2 |
| CSF t-Tau (pg/ml, mean ± SD) | 44 ± 24 | 113 ± 64 | 69 ± 44 |
| CSF p-Tau181 (pg/ml, mean ± SD) | 32 ± 13 | 79 ± 32 | 53 ± 20 |

**Figure 1.**
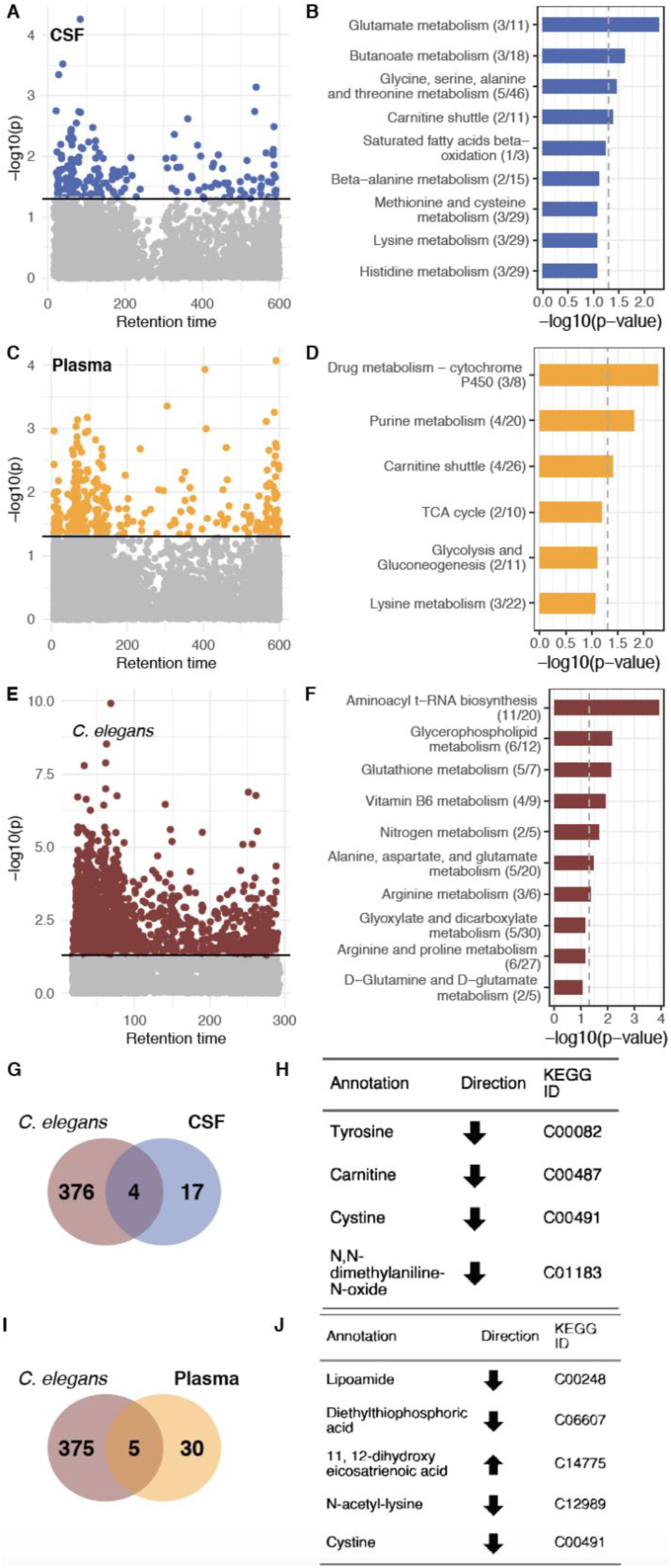
Global metabolomic features associated with p-tau in humans and *C. elegans*. In A, a metabolome-wide association study found 225 CSF metabolites associated with CSF p-tau levels with *p* < 0.05. Enriched pathways corresponding to CSF p-tau associated metabolites are shown in B. In C, 391 plasma metabolites were found to be associated with CSF p-tau levels, with *p* < 0.05; enriched pathways are listed in D. The aggregating strain shows the greatest influence on the metabolome. In E, a metabolome wide association study found 900 metabolites significantly different between the aggregating and non-aggregating worms. These metabolites resulted in the enriched pathways shown in F. In G and I, Venn diagrams shows the overlap between annotated metabolites associated with p-tau in worms and the human matrix; H and J include the name and KEGG ID for the overlapping metabolites. The nature of the relationship between the metabolite and p-tau, whether positively associated (upward arrow) or negatively associated (downward arrow), is shown under the direction column in H and J.

### Changes in global metabolism associated with aggregating tau protein expression in *C. elegans*

HRM detected 19,380 metabolite peaks in *C. elegans* using the HILIC column with positive ionization mode. After blank filtration and imputation, 8,860 were retained for further analysis. Metabolome wide association analysis found more than 900 m/z features that were significantly different (*p* < 0.05) between the aggregating and non-aggregating strain (Figure 1 E). Metabolites were tested for pathway enrichment using the KEGG *C. elegans* reference map, which identified changes in the tryptophan and arginine pathway, glycerophospholipid metabolism, lysine degradation, glutathione metabolism, as well as glutamate and glutamine metabolism implicating altered amino acid metabolism (Figure 1 F).

### Metabolites associated with aggregating tau protein in both species

The analysis to determine metabolites associated with p-tau in both humans and *C. elegans* was conducted separately for CSF and plasma. Metabolite annotations from mummichog (Schymanksi level 3 confidence (Schymanski et al. 2014)) were then used to test for overlap with unique KEGG ID annotation. We identified 4 CSF-derived metabolites and 5 plasma-derived metabolites overlapping with metabolites from the aggregating tau *C. elegans* strain associated with CSF p-tau levels in the same direction (Figure 1 G - J).

### Metabolites associated with AD dementia versus normal controls

Since common thresholds for feature filtering in metabolomics pre-processing pipelines may remove low-abundance exogenous chemicals of interest, we conducted a sensitivity analysis to identify plasma metabolites associated with AD (vs. NC) using a lower threshold for missingness (removal of features missing in >80% of samples) compared to our original study (>20%) (Supplemental Table 2). One feature elevated in AD, *m/z* 386.8946, could not be identified with MS/MS due to low abundance but had a unique match in the METLIN database to 1,1-dichloro-2-(dihydroxy-4’-chlorophenyl)-2-(4’-chlorophenyl)ethylene, a metabolite of DDT.

### DDT uptake in *C. elegans*

The *C. elegans* cuticle is known to be a barrier against absorption of toxicants (Hunt 2017). The nematode is known to possess CYP 450 enzymes, although their repertoire is not as extensive as in mammals (Harlow et al. 2018). We evaluated if *C. elegans* can absorb and metabolize DDT by measuring the levels of p,p’-DDT, p,p’-DDE, p,p’-DDD, o,p’-DDT, o,p’-DDE, and o,p’-DDD in worms using GC-HRMS. In wildtype worms, exposure to 0.3, 3, and 30 μM DDT led to internal levels of 0.27, 0.49 and 1.3 picogram of p,p’-DDT in each worm, respectively (Figure 2 A). All metabolites of p,p’-DDT were also detectable and measured in the transgenic strain. In all strains exposed to 3 μM DDT, the levels of p,p’-DDE were about 5-10 times lower than p,p’-DDT.

**Figure 2.**
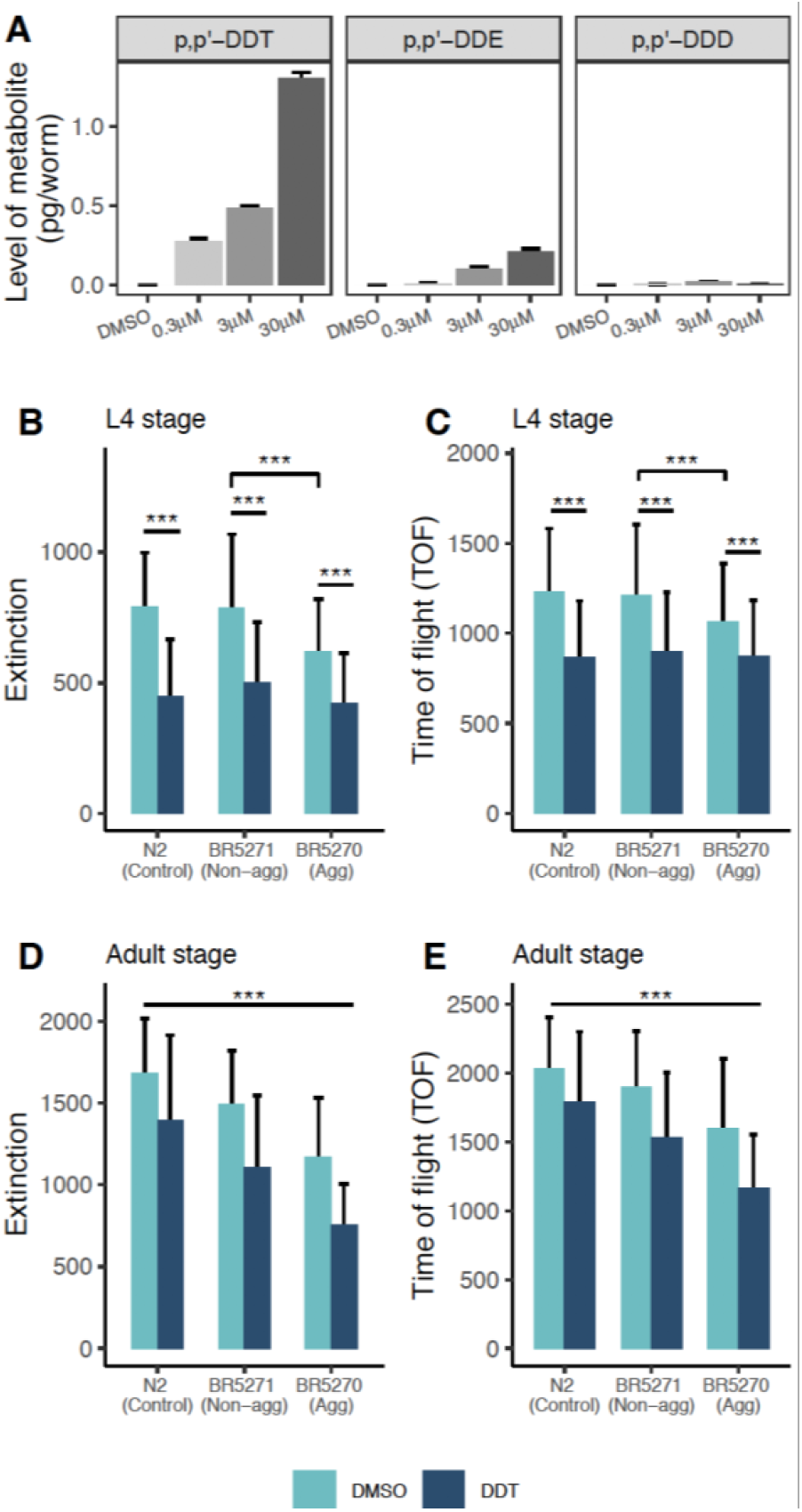
Uptake and metabolism of DDT and the effect of exposure on growth. In wildtype worms, exposure to increasing levels of DDT shows increasing internal levels of p,p’-DDT and its metabolite, p,p’-DDE while levels of p,p’-DDD were near the limit of detection (A). Wildtype worms exposed to the three doses of DDT were collected in triplicate and the mean level of the parent and its metabolites is plotted with error bars representing the standard deviation. The aggregating strain is smaller in size at larval stage 4 (B, C) and in young adulthood (D, E) compared to the non-aggregating and wildtype strain. Exposure to DDT restricted the growth of all strains at both stages measured. The bars represent the mean measure and the error bars represent the standard deviation. *** Tukey HSD adjusted p < 0.0001.

### Effect of DDT exposure on *C. elegans* size

Assessment of TOF (length) and extinction (optical density) at 46-50 hours post synchronization (∼larval stage 4) revealed that the aggregating strain were smaller and with lower density compared to the non-aggregating and wildtype worms. Exposure to DDT reduced the size and density in all strains assessed (Figure 2 B & C). Measurement at 70-72 hours post synchronization (young adulthood) showed that both tau transgenic strains are smaller (p < 0.0001) than wildtype worms, with the aggregating strains more severely affected. Exposure to DDT reduced the size of all strains in a graded manner, with the aggregating strain exposed to DDT being the smallest (Figure 2 D & E).

### Effect of tau protein and DDT on swim behavior of *C. elegans*

The aggregating strain showed differences in wave initiation rate and travel speed compared to the wildtype worms (Figure 3 A & B). Exposure to DDT in the aggregating strain almost doubled the percentage of time the worms spent curling when swimming (average percentage of time spent curling in aggregating worms + solvent control was 1.13%, and in aggregating worms + DDT: 1.83%, p < 0.001, Figure 3 C). The aggregating strain showed a lower activity index (average activity index in non-aggregating strain was 388.6 and in aggregating strain was 286.6, p < 0.01) Figure 3 D), and a difference in time spent reversing and brushstroke (p < 0.05 Supplemental Figure 1) compared to wildtype worms. Exposure to DDT did not significantly alter any swim behavior in the wildtype or non-aggregating strain.

**Figure 3.**
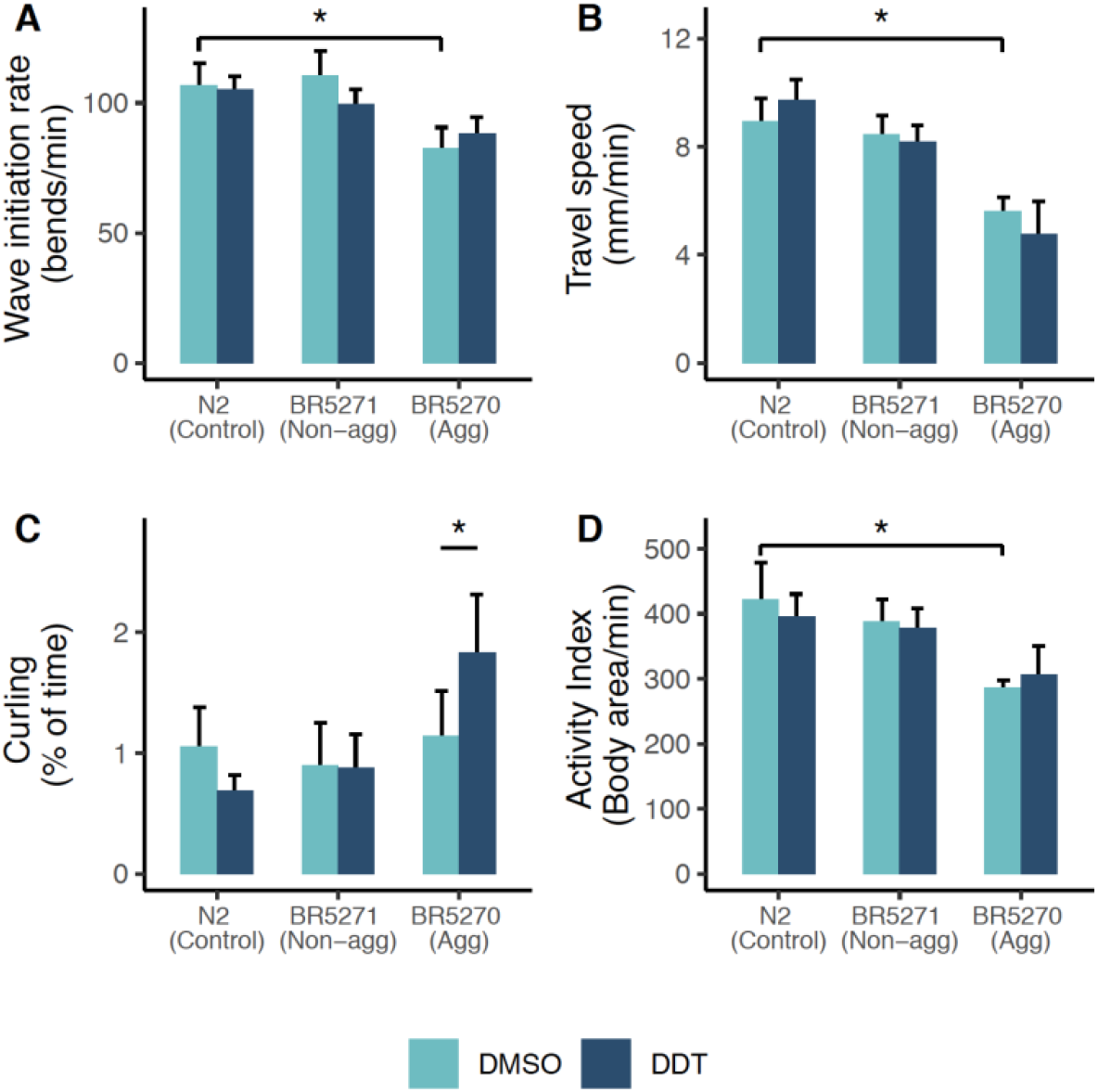
Aggregating tau and DDT affect swimming behavior. The aggregating strain shows altered wave initiation rate (A) and travel speed (B) compared to the wildtype N2 worm. The aggregating strain exposed to DDT spends more time curling while swimming (C). The overall activity index of the aggregating strain is reduced compared to the wildtype N2 strain (D). Each bar represents the mean measure taken from four-five different trials with a total of 50-100 worms per group. The error bars represent the standard error of the mean. * Tukey HSD adjusted *p* < 0.05.

### Effect of tau and DDT on mitochondrial respiration

Wildtype worms and the non-aggregating strain showed similar oxygen consumption profiles (Figure 4 A). The aggregating strain showed reduced rates of basal, maximal, spare, and non-mitochondrial oxygen consumption rate (OCR) when compared to the non-aggregating strain (Figure 4 B - E). Exposure to 3 μM DDT in the N2 and non-aggregating strain reduced OCR at all four states (Figure 4 A - E). Exposure to DDT in the aggregating strain significantly reduced basal OCR (p < 0.05, Figure 4 B). Other measures of respiration were also reduced due to DDT exposure in the aggregating strain however, none were significantly different at p < 0.05 (Figure 4 C - E).

**Figure 4.**
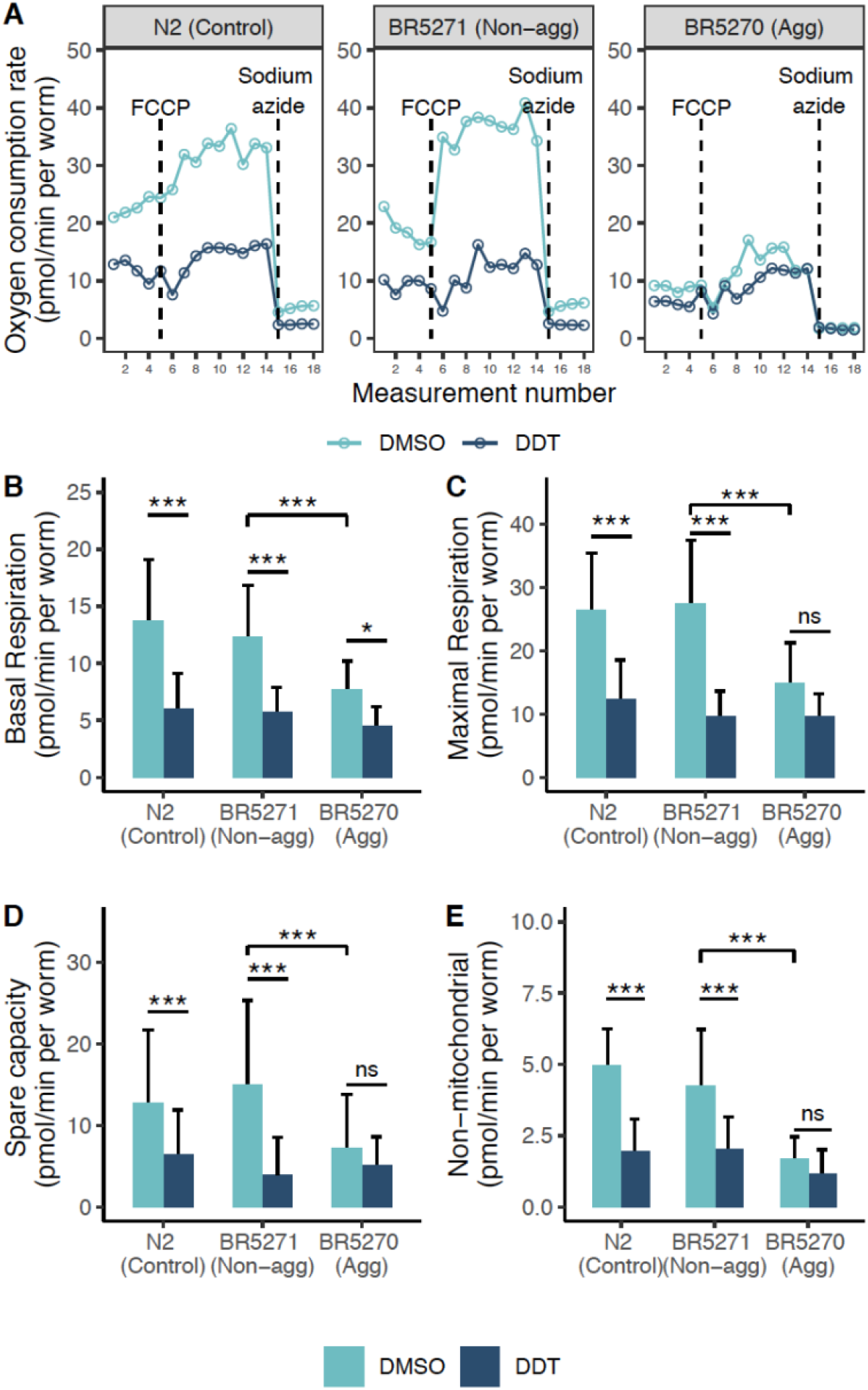
Aggregating tau and DDT inhibits mitochondrial function. In A, a representative oxygen consumption rate (OCR) profile measured using the Seahorse respiratory flux analyzer. The wildtype and non-aggregating strain show a similar OCR however, exposure to DDT reduced the OCR in both strains (A). The aggregating strain shows a reduced OCR compared to the wildtype and non-aggregating strain (A) and exposure to DDT reduced basal respiration in the aggregating strain (B). Each bar represents the mean respiratory measure made across 3-5 experiments with 7-12 wells per run with 3-30 worms per well. The error bar represents the standard deviation. *** Tukey HSD adjusted p < 0.0001, * Tukey HSD adjusted *p* < 0.05, ns: not significant.

### Metabolic response to DDT and tau protein

All strains exposed to DDT showed lower metabolite intensities using HILIC with positive ionization (Figure 5 A) and the other modes: HILIC column under negative ESI and the C18 column under positive and negative ESI (Supplemental Figure 2 - 4). A biplot of principal component (PC) 1 against PC2 shows that the wildtype and non-aggregating strain cluster together while the strains exposed to DDT clustered differently from the unexposed wildtype and non-aggregating strain along PC1 (Figure 5 B). Levels of uric acid and adenosylselenohomocysteine were elevated in all strains exposed to DDT (Figure 5 C). In wildtype worms, exposure to DDT altered amino acids pathways (Figure 6). DDT exposure in the aggregating strain resulted in altered amino acid pathways, TCA cycle metabolism and glyoxylate and dicarboxylate metabolism (Figure 6).

**Figure 5.**
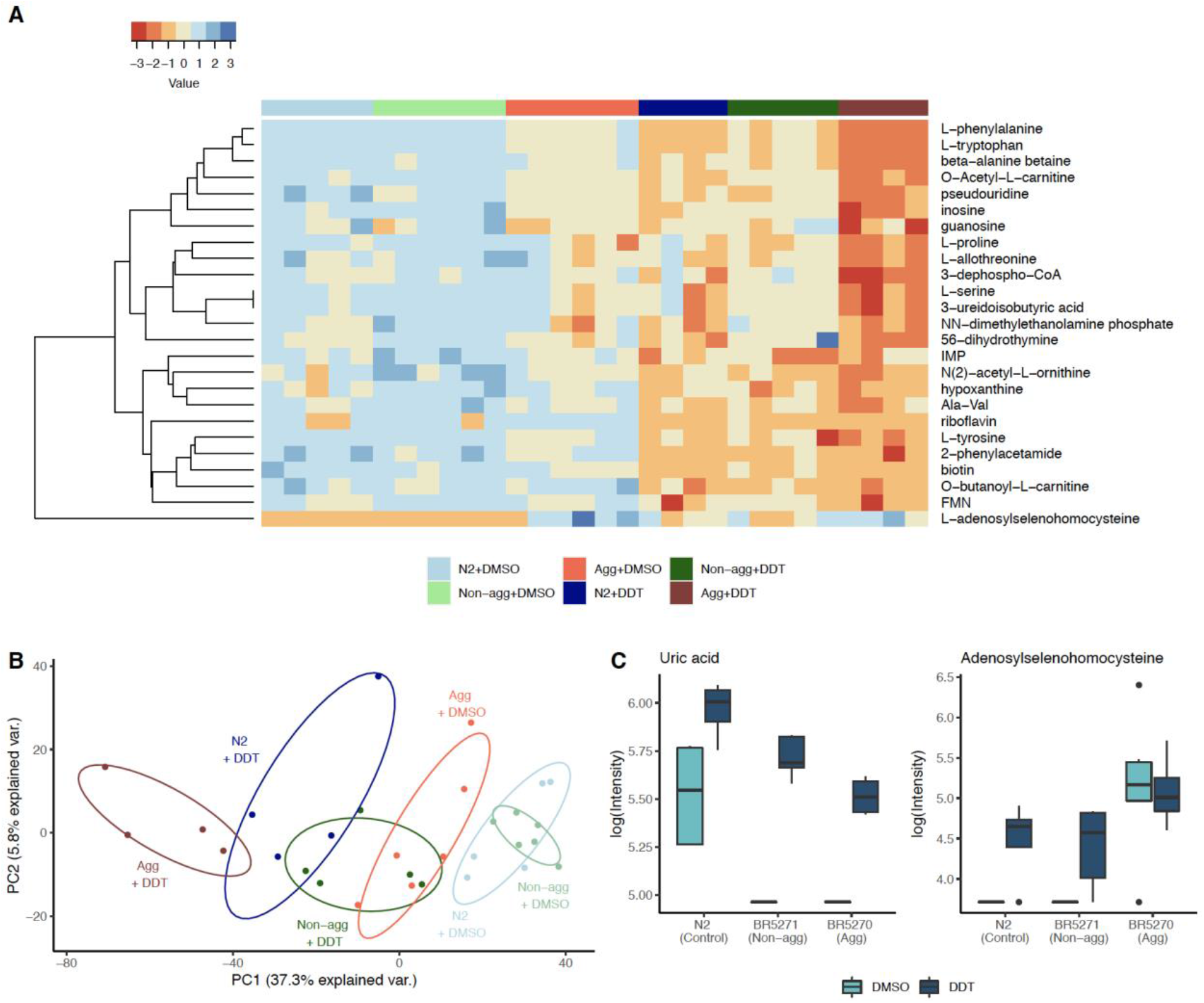
Metabolomic profile of DDT exposure. In A, a heatmap of the top 25 metabolites with the smallest *p-value* hierarchically clustered shows that in all strains, DDT decreases metabolite intensity. In B, a PCA biplot of PC1 plotted against PC2 shows that strains exposed to DDT cluster differently from the wildtype and non-aggregating control strains. The aggregating strain does not cluster with the wildtype and non-aggregating control groups, suggesting variation as a result of aggregating tau protein expression. In C, levels of uric acid are higher in all worms exposed to DDT and levels of adenosylselenohomocysteine are higher in both: worms exposed to DDT and in worms expressing aggregating tau protein. IMP: inosine monophosphate, FMN: flavin mononucleotide, Ala-Val: alanine-valine dipeptide.

**Figure 6.**
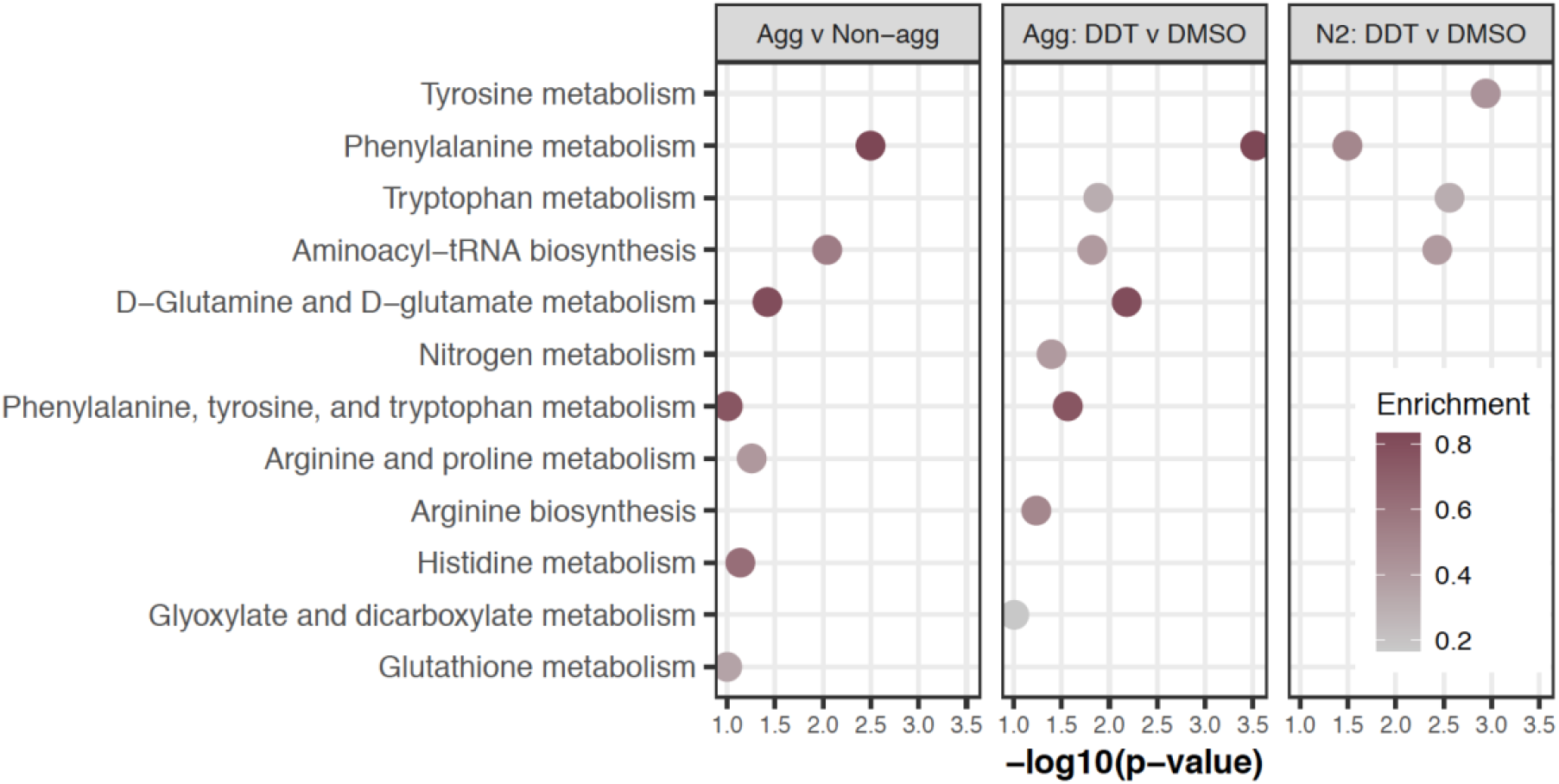
Pathway analysis. The different metabolic pathways enriched in the three different comparison groups. Enrichment is calculated as the ratio of the number of significant hits to the total pathway size.

### Effect of DDT exposure on learning

There was no difference in learning determined through the associative learning paradigm among the three strains. Further, exposure to DDT did not show an effect on learning (Figure 7 A & B).

**Figure 7.**
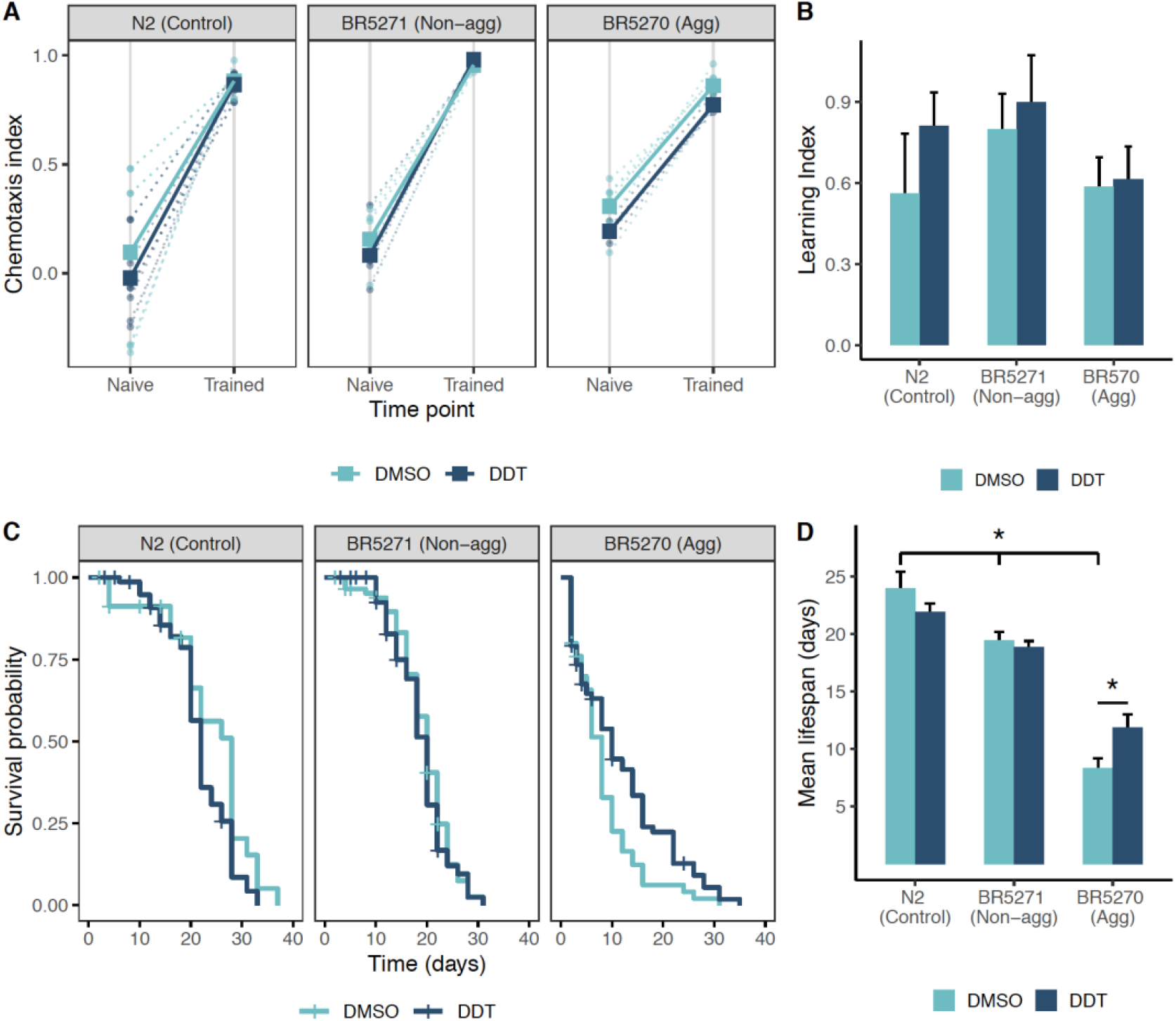
Associative learning and survival. Associative learning was not affected by exposure to DDT in any of the strains. The aggregating strain did not learn differently from the non-aggregating or wildtype worms. The dotted lines represent the chemotaxis index for each trial with the bold lines representing the mean (A). Each bar represents the learning index (B), calculated as the difference between the trained chemotaxis index, post-conditioning, and the naïve chemotaxis index for each trial (A). The error bars represent the standard error of the mean. The non-aggregating and aggregating strain live shorter than wildtype worms. Exposure to DDT did not affect the survival of the wildtype or non-aggregating strains however, exposure to DDT slightly rescued the reduced lifespan in the aggregating strain. The Kaplan Meier curves (C) are generated by following 60 – 120 worms in each group and the bars (D) represent the mean lifespan in days and the error bars represent the standard deviation. * Tukey HSD adjusted *p* < 0.05.

### Effect of DDT exposure on survival

The non-aggregating and aggregating strains exhibited a shorter lifespan compared to the wildtype worms (average lifespan in wildtype worms was 24 days, in non-aggregating worms was 19.4 days, and in aggregating worms was 8 days, p < 0.0001, Figure 7 C). Exposure to DDT does not alter the lifespan in wildtype or the non-aggregating strain. In the aggregating strain, exposure to DDT slightly rescued the reduction in lifespan in the strain (mean lifespan was 11.8 days), however, the lifespan was still shorter than that of the non-aggregating and wildtype strain (Figure 7 C & D).

## Discussion

Model organisms are a useful tool to understand age-related changes in biology and pathology. A number of signaling pathways that act as master regulators of lifespan are conserved in yeast, nematodes, flies, and mammals (Bishop et al. 2010). The use of model systems has uncovered evolutionarily conserved pathways that regulate both longevity and age-related changes in learning and memory (Bishop et al. 2010; Friedman and Johnson 1988; Grotewiel et al. 2005; Kauffman et al. 2010; Kenyon et al. 1993; Klass 1977; Silva et al. 1998). We used the nematode *C. elegans* to study environmental determinants of aging and cognitive function. The organism’s short lifespan (2-3 weeks) makes it ideal to study the process of aging and diseases associated with age, such as AD (Ardiel and Rankin 2010; Arey and Murphy 2017; Jonsson et al. 2013; Link 2006). In addition, *C. elegans* mitochondria show close structural and functional conservation to mammalian mitochondria (Murfitt et al. 1976) and pathways of intermediary metabolism are also highly conserved (O’Riordan and Burnell 1990). Thus, we attempted to find similarities in systemic biochemistry associated with aggregating tau protein toxicity, which is a pathological hallmark of AD, in humans and *C. elegans*.

Our group has previously reported plasma metabolites associated with AD in the cohort studied herein (Niedzwiecki et al. 2020). The most significant (lowest p-value) metabolite associated with CSF p-tau was a metabolite of the drug Rivastigmine, an acetylcholinesterase inhibitor. This was also the top metabolite associated with AD in the previous study. The plasma levels of glutamine were positively associated with levels of CSF p-tau. Metabolomic profiling of the CSF showed a negative association between CSF levels of p-tau and glutamine, contrary to the direction of the association found in plasma. This could be the result of differential changes in the glutamate/glutamine cycle in the central nervous system and the periphery. The CSF-derived metabolites associated with p-tau levels are related to butanoate metabolism and carnitine shuttle pathways, both of which are associated with mitochondrial function (Pettegrew et al. 2000; Rose et al. 2018).

Using a transgenic strain of *C. elegans* that expresses a mutant form of human tau protein in all neurons, we observed changes in several metabolite peaks that were associated with aggregating tau protein. Pathway analysis using these features revealed changes in metabolic pathways that have previously been associated with neurodegeneration and AD, including the glycerophospholipid pathway, and glyoxylate and dicarboxylate metabolism (Frisardi et al. 2011; Yan et al. 2020)

The 4 metabolites associated with p-tau and overlapping between CSF metabolites and worm metabolites were tyrosine, carnitine, cystine and N, N-dimethylaniline N oxide. While we did not find information on the relationship between N, N-dimethylaniline N oxide and neurodegeneration or AD, all three of the other metabolites have been previously associated with AD. Several studies have reported lower levels of tyrosine measured in the CSF of AD patients (Basun et al. 1990; Martinez et al. 1993). An untargeted HRM analysis of CSF samples from MCI patients showed altered tyrosine metabolism (Hajjar et al. 2020). A study of CSF from non-*APOE4* carriers in the early stage of AD reported lower levels of carnitine in the CSF (Lodeiro et al. 2014). Another study of CSF from AD patients found lower levels of free carnitine but increased levels of acylcarnitine suggesting impaired energy production through anaplerotic pathways (van der Velpen et al. 2019). We detected decreased levels of cystine in plasma, CSF, and worms. Cystine is the dimer form of cysteine, a sulfur containing amino acid that functions to reduce redox stress. Several studies have reported increased levels of cysteine in the brain, plasma, and CSF of AD patients (Czech et al. 2012; Mahajan et al. 2020; Trushina et al. 2013). This could be a result of increased conversion of cystine to cysteine to ameliorate oxidative stress.

Apart from cystine, the other metabolites associated with p-tau found to be overlapping between plasma and worms were lipoamide, diethylthiophosphoric acid, 11,12-dihydroxy-5Z,8Z,14Z-eicosatrienoic acid (11,12-DHET), and N-acetyl lysine. Lipoamide is the amide form of lipoic acid which is a naturally occurring disulfide compound that functions as a co-factor for mitochondrial bioenergetic enzymes. It has been proposed as a novel treatment for AD owing to the many functions it performs (Holmquist et al. 2007). Lipoic acid can increase acetylcholine production (Haugaard et al. 2000) and glucose uptake (Holmquist et al. 2007) and it is reported to improve peripheral insulin resistance and impaired glucose metabolism (Bitar et al. 2004; Lee et al. 2005; Thirunavukkarasu et al. 2004). Diethylthiophosphoric acid is a part of the aminobenzoate degradation pathway. Derivatives of aminobenzoic acid may have potential as drugs to inhibit acetylcholinesterase, thereby ameliorating the acetylcholine deficit present in AD (Shrivastava et al. 2019). 11,12-DHET derives from oxidation of arachidonic acid, a well-known precursor activated during inflammatory response. A study using strains of mice expressing Aβ and tau in the brain found increased levels of several eicosanoids in the brain and in plasma of these mice (Tajima et al. 2013). In both human plasma and worms we found higher levels of DHET, in line with previous findings. Finally, altered lysine metabolism has been previously reported in cases of MCI compared to cognitively normal individuals (Trushina et al. 2013) and lysine supplementation has been proposed as a treatment strategy for AD (Kumar and Kumar 2019).

Neither the human nor *C. elegans* metabolome is fully curated, and non-targeted metabolomics data includes many dietary, microbiome and environmental chemicals in addition to those associated with endogenous metabolic pathways as presented here. Although the results from this cross-species analysis should be interpreted with caution, the concordance between several metabolites that have been previously associated with AD provides support for using *C. elegans* as a model to study biochemical changes associated with AD-related pathology. Further, the disruptions in evolutionarily conserved pathways that are associated with AD-related related pathology offer great power for mechanistic interpretation. Correlation between metabolites observed across species could provide a means to identify overlapping central networks and interacting sub-networks associated with AD-related pathology (Kalia et al. 2019). In the future, we plan to use mutant strains and appropriate exposures to determine the role of these metabolites in the aggregating tau protein related toxicity in *C. elegans*.

Untargeted HRM approaches allow us to study the effect of the exposome on human health (Vermeulen et al. 2020). However, in untargeted HRM analyses, the abundance of exogenously-derived parent compounds and their metabolites tend to be orders of magnitude lower than endogenous chemicals (Rappaport et al. 2014) and may not be present in all study participants. Thus, statistical approaches and thresholds need to be adjusted to account for this lower abundance and prevalence of exogenous chemicals in population studies. Therefore, we conducted a sensitivity analysis by applying a lower threshold for feature filtering in our previous analysis of plasma-derived features associated with AD. The analysis found higher levels of a halogenated metabolite in the plasma of AD patients, which was putatively identified as a derivative of the persistent pesticide DDT (Supplemental Table 2).

To investigate whether exposure to DDT can exacerbate tau protein toxicity, we used a transgenic *C. elegans* strain that expresses human tau protein and a mutated tau protein sequence that has a propensity to form tau protein aggregates. We also used a transgenic strain to serve as control for the aggregating strain that expresses the same human tau protein but the mutated tau protein sequence is not prone to aggregation. These transgenes are expressed in all neurons of the worm driven through the *rab-3* promotor (Fatouros et al. 2012).

The targeted GC-HRMS assay detected and measured several metabolites of p,p’-DDT in worms exposed to the pesticide, suggesting that the pesticide is not only absorbed but also biotransformed in the nematode, supporting the use of this model to study the toxic effects of DDT. Previously, Mahmood (2016) found that exposure to 1 μg/mL of DDT (∼2.8 μM) had a mild inhibitory effect on pharyngeal pumping, while this dose had no effect on brood size. A survey of serum samples analyzed for levels of p,p’-DDT conducted by NHANES showed a wide range of the pesticide in the blood of the American population, and levels of p,p’-DDT measured increased with increasing age. Among those aged 12-19 years in the survey, the geometric mean of lipid adjusted serum p,p’-DDT level was less than 5 ng/g lipid (CDC 2020). Assuming the wet mass of a single worm is 1 μg (Muschiol et al. 2009), exposure to 3 μM DDT using our paradigm resulted in a mean level of ∼0.5 ng/g wet weight of *C. elegans*. Thus, we exposed the wildtype, aggregating, and non-aggregating strains to 3 μM DDT during development and measured the effect of exposure on swim behavior, respiration, growth, metabolism, learning, and lifespan.

Similar to previous findings (Fatouros et al. 2012), we observed that the aggregating strain travels slower than the wildtype worms. The aggregating strain also showed a reduced wave initiation rate, which is akin to a swimming stroke rate (Restif et al. 2014), compared with the non-aggregating and wildtype strain. Additionally, the aggregating strain had a lower overall activity index compared with the non-aggregating and wildtype strain. Exposure to DDT significantly increased the amount of time the aggregating strain spent curling. The curling phenotype has been used to screen for motility defects in worms. A recent screen for curling identified the *bcat-1* gene to be associated with a Parkinson’s-like phenotype and knockdown of the gene transcript showed altered mitochondrial function (Mor et al. 2020). The curling phenotype has also been used to ascertain dopaminergic toxicity due to the complex I inhibitor, MPP^+^ (Braungart et al. 2004; Richardson et al. 2005).

We observed that the aggregating strain has severely impaired mitochondrial respiration, with diminished basal and maximal respiration compared with the non-aggregating strain. Several *in vitro* and *in vivo* studies have shown that aggregating tau protein can inhibit complex I and V of the mitochondria (David et al. 2005; Kim and Chan 2001; Lasagna-Reeves et al. 2011). Tau protein can alter the mitochondrial membrane potential, cause activation of the apoptotic-related caspase-9, and impede energy production (Lasagna-Reeves et al. 2011; Shafiei et al. 2017). Furthermore, disintegration of tau protein can lead to disturbed transport of mitochondria across microtubules and mitochondrial fission-fusion dynamics (Eckert et al. 2014; Fatouros et al. 2012).

Exposure to DDT in the wildtype and non-aggregating strain severely impaired mitochondrial respiration at baseline and in the uncoupled state (FCCP). Several *in vitro* and *in vivo* studies have reported an inhibitory effect of DDT on mitochondrial function and ATP production, but none have reported this in *C. elegans*. DDT is known to inhibit complex II, III, and V of the electron transport chain and it depresses the mitochondrial membrane potential (Elmore and La Merrill 2019; Moreno and Madeira 1991). In rats, exposure to DDT reduced the number of mitochondria measured in the liver and altered fatty acid metabolism (Liu et al. 2017), an effect that would be consistent with the overall decrease in OCR under all states measured. We note that this overall decrease in oxidative phosphorylation activity is also unlikely to be attributable to decreased motility since exposure to DDT did not affect the swimming behavior of wildtype worms, reinforcing the interpretation that DDT may directly affect either mitochondrial content and/or respiratory chain activity.

We found that the aggregating strain was smaller in size at larval stage 4 and in adulthood compared to the non-aggregating strain. This could be attributed to reduced energy production and biomass accumulation due to mitochondrial inhibition. When exposed to DDT, there was no difference in size between the aggregating and non-aggregating strains at larval stage 4; however, in adulthood, DDT reduced the size of the aggregating strain more than in the non-aggregating strain. It is likely that the additional insult on the mitochondria by DDT restricts growth which was only detectable following the initiation of reproductive capacity.

HRM found several metabolites to be altered in worms following DDT exposure. Levels of several amino acids were reduced along with intermediates of the TCA cycle and branched chain amino acid metabolism. Uric acid was increased in all strains exposed to DDT (Figure 6 A). Uric acid is the end product of purine metabolism and has anti-oxidant properties since it can scavenge free radicals and prevent lipid peroxidation (Hooper et al. 1998). High levels of uric acid have been reported to induce stress response pathways in *C. elegans* by increasing levels of the DAF-16/FOXO and SKN-1/NRF-2 transcripts (Wan et al. 2020). Levels of adenosylselenohomocysteine were also found to be increased in all strains exposed to DDT and in the aggregating strain (Figure 6 B). Thioredoxin reductase-1 (TrxR-1) is the only selenium containing protein in *C. elegans* (Rohn et al. 2018). An elevated seleno-metabolite suggests increased levels of TrxR-1 in response to oxidative stress induced by DDT exposure and tau protein aggregation.

The aggregating strain did not show any difference in their ability to learn following an associative training paradigm, compared to the non-aggregating or wildtype strain. These findings are similar to those made by Wang and colleagues (2018). Exposure to DDT did not affect this ability to learn in either strain using the associative learning assay. We found that the non-aggregating and aggregating strains have a reduced lifespan compared to wildtype worms, replicating previous findings (Wang et al. 2018). The proteotoxicity and reduced respiratory rate in the aggregating strain could explain this observation (Zarse et al. 2007). Interestingly, exposure to DDT did not change the mean lifespan in wildtype or non-aggregating worms but it slightly increased the mean lifespan of the aggregating strain. This finding is surprising but given that the exposure occurred developmentally, it hints to the activation of mitohormetic pathways which could turn on lifespan extension pathways (Maglioni et al. 2019), like the mitochondrial unfolded protein response (UPR^mt^) pathway. However, the extension in lifespan was not large enough to be as much or more than the lifespan of the non-aggregating or wildtype strain. It is also possible that, while mitochondrial inhibition by aggregating tau protein alone does not induce the UPR^mt^ pathways, the mitochondrial stress induced by DDT during development produces an antagonistic effect which induces stress response pathways (Wytock et al. 2020).

While we present evidence that supports the use of *C. elegans* as a model to study whether DDT can exacerbate tau protein toxicity, our study has several limitations. In insects and mammals, DDT inhibits voltage-gated sodium channel inactivation and stabilizes the open state of sodium channels, causing prolonged channel opening (Bloomquist 1996; Narahashi 2000). The *C. elegans* genome does not encode for voltage-gated sodium channels (Hobert 2018), thus DDT does not produce neurotoxicity through this mechanism in the nematode. Thus, we were unable to measure any interaction tau protein aggregation may have with altered neuronal excitability elicited by DDT in mammalian neurons. Furthermore, in the transgenic model we chose, we were unable to control the level of tau protein aggregates expressed in the neurons. It is possible that the severe tau protein aggregation toxicity obscured effects of DDT exposure and its proteotoxic effects. The interactions between the two insults may become more apparent when lower levels of the aggregates are expressed.

Despite these limitations, we provide evidence that support the use of *C. elegans* as a model to study gene-environment interactions. We provide evidence that DDT is taken up and biotransformed by *C. elegans*. In wildtype worms, DDT restricts growth, as measured by size, and reduces mitochondrial respiration. DDT produces major changes in global metabolism, including pathways related to neurotransmitter precursors and other amino acid metabolism. In transgenic worms that express an aggregating form of human tau protein in all neurons, DDT restricts growth even further and reduces the basal respiration rate. Aggregating tau worms exposed to DDT spend more time curling when swimming, a known mitochondrial toxicity phenotype. Further, DDT exposure affects the metabolism of several amino acids, the TCA cycle, and the glyoxylate and dicarboxylate metabolic pathway. Our data suggest that exposure to DDT likely exacerbates the mitochondrial inhibitory effects of aggregating tau protein in *C. elegans*. Additionally, the concordance between several metabolites that have been previously associated with AD provides validity to using *C. elegans* as a model to study biochemical changes associated with AD-related pathology. In the future, using transgenic *C. elegans* strains, we will perform systematic analyses of the environmental drivers of AD that can lead to interventional strategies aimed at preventing or treating the disease.

## Acknowledgements

G.W.M is supported by NIH grants RF1AG066107, R01AG067501, U2C ES030163, R01ES023839, M.L.B. is supported by NIEHS grant T32ES007732, D.I.W. is supported by NIEHS grant U2C ES030859 and K.E.M. is supported by NEIHS grant T32ES007272.

## Data sharing

The patient related metabolomics data will be made available on metabolomics workbench. All data and code related to DDT exposure in *C. elegans* is available through a repository on VK’s github account https://github.com/vrindakalia/DDT_tau_Celegans.

