## supplemental_tables_and_figures for "Cross-species metabolomic analysis of DDT and Alzheimer’s disease-associated tau toxicity"

**Supplemental Table 1.** Retention time and ions monitored to quantify and confirm DDT metabolites

| Analyte | Retention Time (min) | Quantifying Ion (m/z) | Confirming Ion 1 (m/z) | Confirming Ion 2 (m/z) | LOD (ppb) |
|---|---|---|---|---|---|
| o,p'-DDE | 11.13 | 245.9999 | 247.9968 | 317.9345 | 0.016 |
| o,p'-DDT | 12.80 | 235.0076 | 165.0699 | 237.0047 | 0.010 |
| p,p'-DDE | 11.70 | 245.9999 | 247.9968 | 317.9345 | 0.029 |
| p,p'-DDT | 13.32 | 235.0076 | 165.0699 | 237.0047 | 0.029 |
| 4,4'-DDE ($^{13}C_{12}$) | 11.70 | 260.0370 | 188.1021 | 258.0400 | |
| 4,4'-DDT ($D_8$) | 13.27 | 243.0576 | 173.1200 | 245.0549 | |

**Supplemental Table 2.** Lowering the filtering threshold revealed a positive association between a putative DDT metabolite and risk of Alzheimer's disease (AD). [a]ID level indicates annotation confidence: 1, m/z and retention time confirmed with MS2, 2: Multiple/isotopes present; 3: m/z matched single adduct mass within 10 ppm mass error, 4: m/z matched adduct mass of multiple isobaric species, probable identifications listed. RT: retention time.

| m/z | RT | Change in AD | Putative compound(s) | Predicted adduct | ID levels[a] | Notes |
|---|---|---|---|---|---|---|
| 129.0661 | 89 | Higher | Glutamine (2 ppm) | $-H_2O+H$ | 1 | -- |
| 231.1205 | 211 | Higher | 5S,6S-epoxy-15R-hydroxy-ETE-(+Na, 0 ppm) | -- | 3 | -- |
| 246.9550 | 127 | Higher | Numerous database matches | $-H_2O+H$ | -- | Contains halogen (Cl and/or Br) |
| 334.1410 | 86 | Lower | Piperettine (1 ppm) | +Na | 4 | -- |
| 349.1515 | 80 | Lower | Piperine (1 ppm) | +ACN+Na | 4 | -- |
| 386.8946 | 61 | Higher | 1,1-Dichloro-2-(dihydroxy-4'-chlorophenyl)-2-(4'-chlorophenyl)ethylene (9 ppm) | +K | 2 | Contains halogen (Cl and/or Br) |
| 662.0933 | 158 | Higher | GDP-D-mannuronate (+ACN+H[M+1], 0ppm); Chaetocin (-2H2O+H[M+1], 8 ppm); Blighinone (+H[M+1], 9 ppm) | [M+1] isotope | 4 | -- |
| 663.4524 | 36 | Higher | Lipid A-disaccharide-1-P(+2H, 2 ppm); Aluminium dodecanoate (+k, 2 ppm) | -- | 4 | -- |

**Supplemental Figure 1.** Six different swim behaviors measured in wildtype, aggregating, and non-aggregating worms exposed to DDT including asymmetry, attenuation, brush stroke, reverse swim, stretch and body wave number. The aggregating strain spent more time swimming in reverse than the other strains or treatments. The aggregating strain also has a lower brush stroke compared to the wildtype worms. None of the other swim behaviors were significantly different between the different strains or treatments.

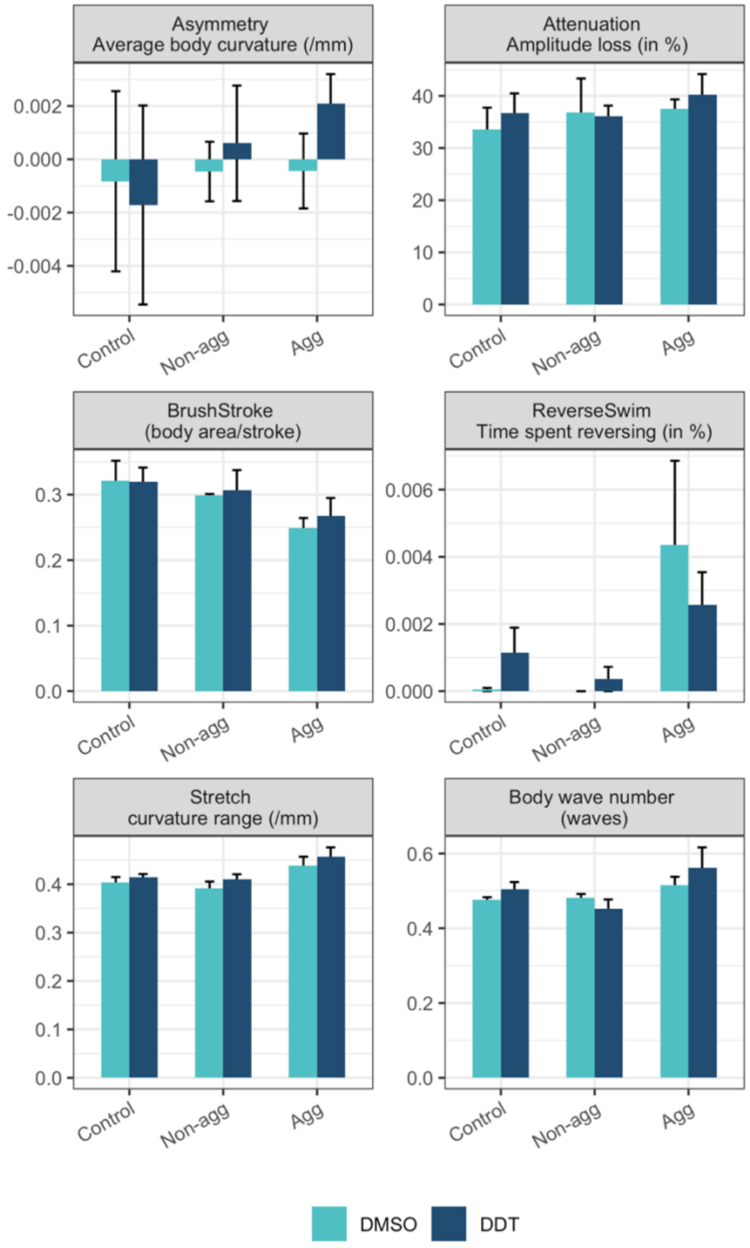

**Supplemental Figure 2.** The heatmap shows annotations and clustering of features measured using the HILIC column under negative electron spray ionization.

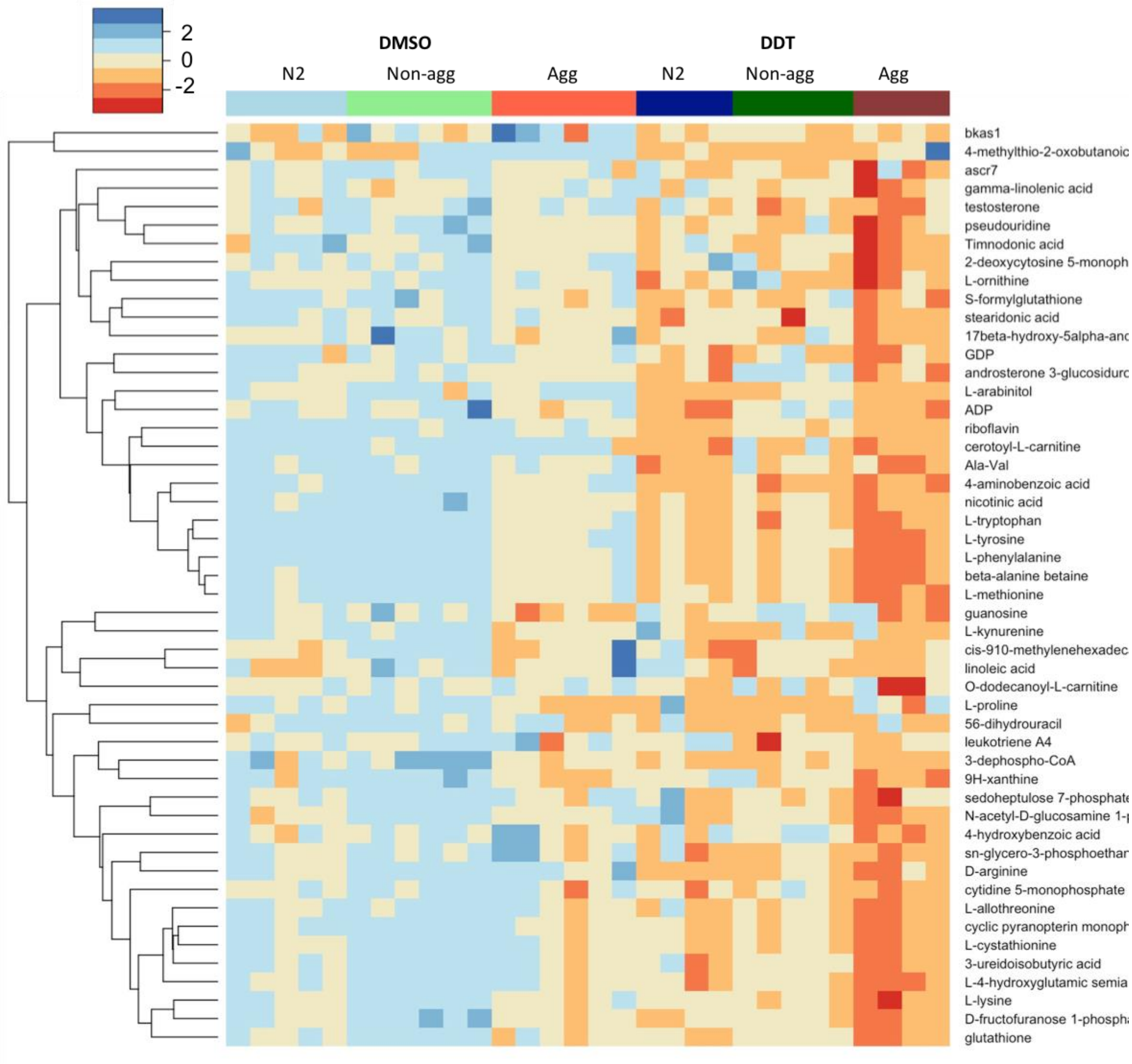

**Supplemental Figure 3.** The heatmap shows annotations and clustering of features measured using the C18 column under positive electron spray ionization.

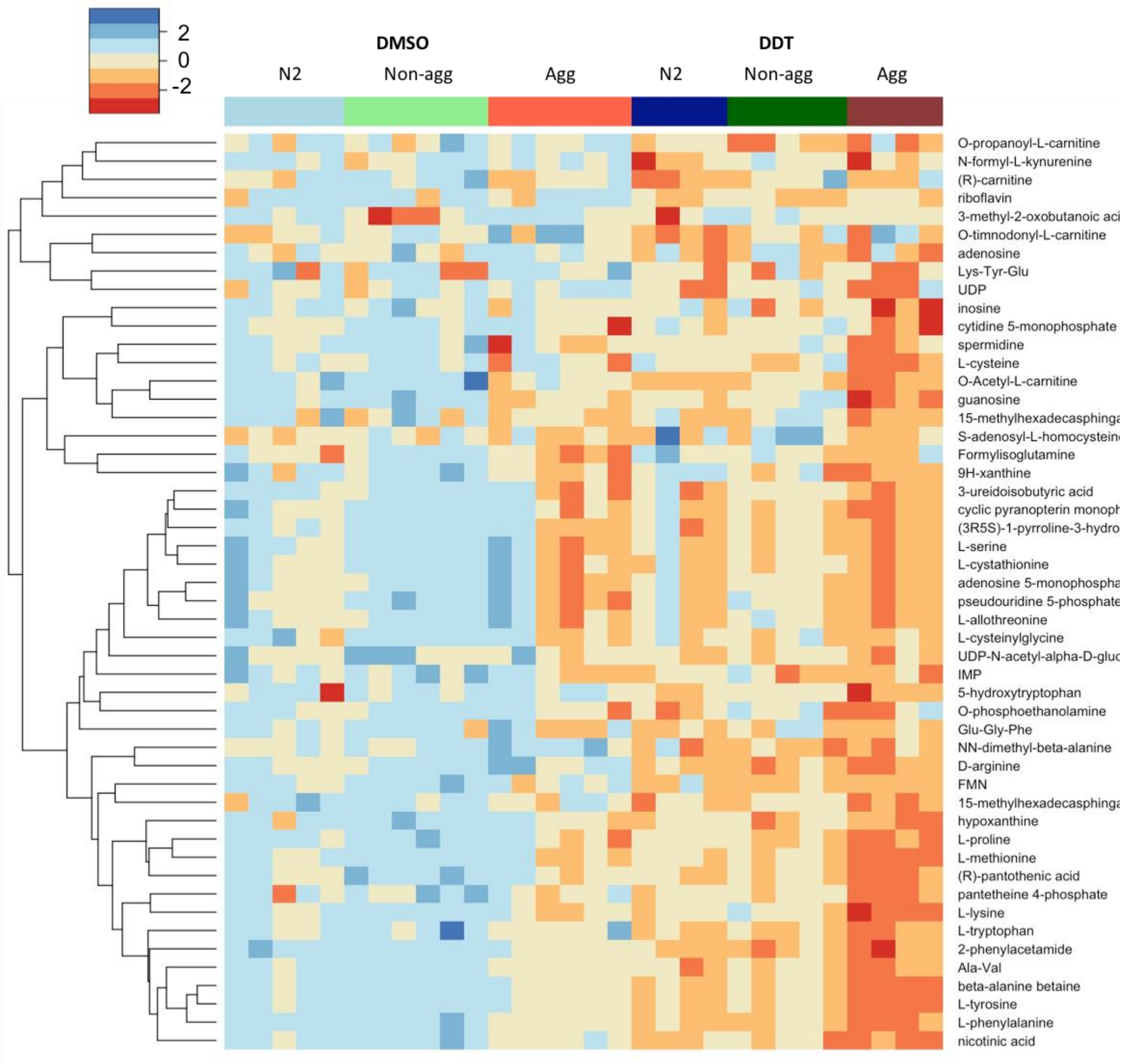

**Supplemental Figure 4.** The heatmap shows annotations and clustering of features measured using the C18 column under negative electron spray ionization.

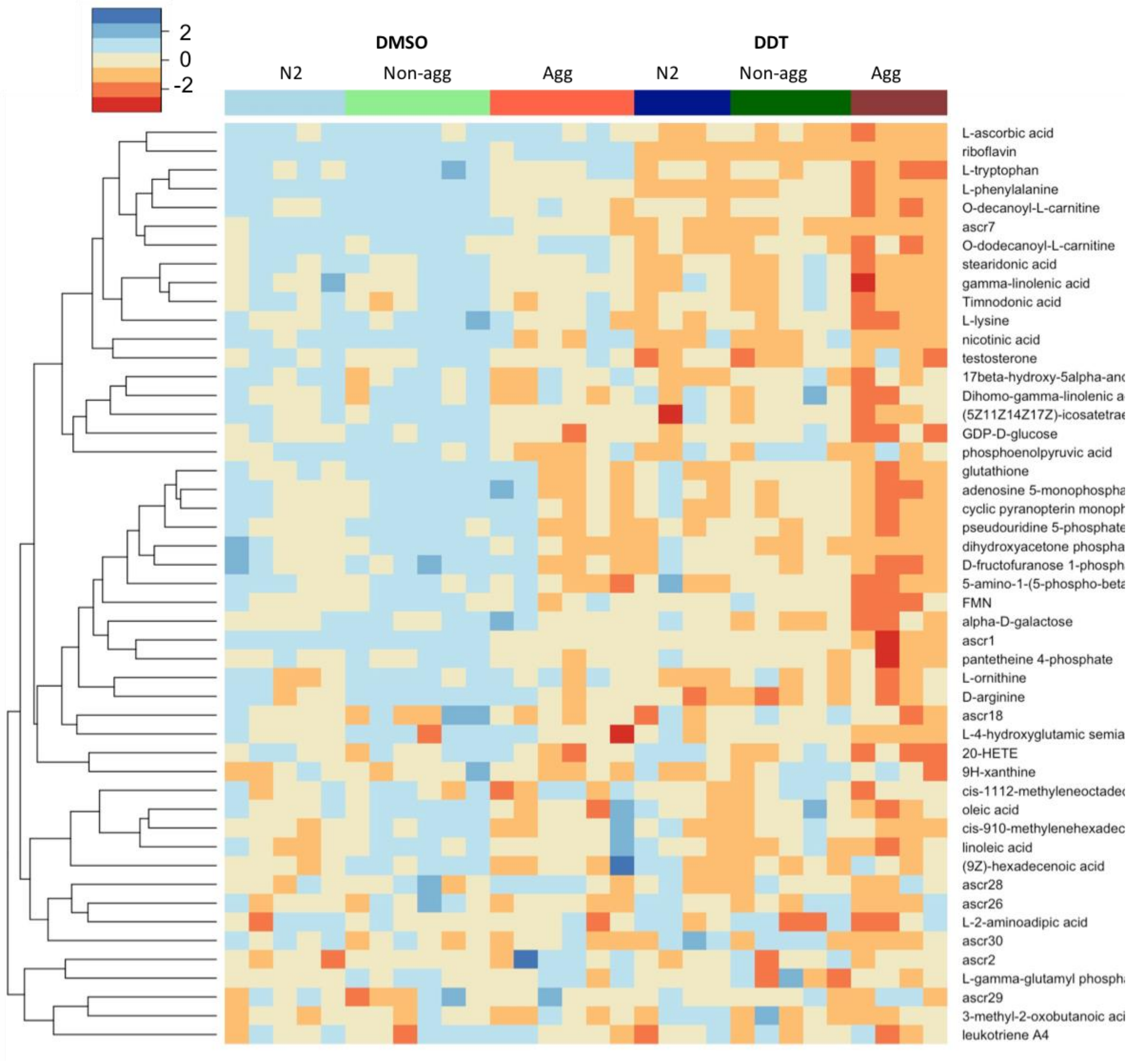
